# Comparative analysis of intracellular and extracellular antibiotic resistance gene abundance in anaerobic membrane bioreactor effluent

**DOI:** 10.1101/702076

**Authors:** Phillip Wang, Moustapha Harb, Ali Zarei-Baygi, Lauren B. Stadler, Adam L. Smith

**Affiliations:** Astani Department of Civil and Environmental Engineering, University of Southern California, 3620 S Vermont Ave, Los Angeles, CA 90089, USA; Department of Civil and Environmental Engineering, Rice University, 6100 Main Street, Houston, TX 77005, USA

**Author notes:** Corresponding author (Adam L. Smith).

## Abstract

The growing practice of wastewater reuse poses a significant risk to further dissemination of antibiotic resistance due to the abundance of antibiotic resistance bacteria (ARB) and antibiotic resistance genes (ARGs) in wastewater effluents. Anaerobic membrane bioreactors (AnMBRs) are an emerging wastewater treatment technology capable of reducing the total ARGs and ARB load discharged to receiving environments compared to conventional aerobic treatment processes. While size exclusion is effective at retaining ARB and its associated intracellular ARGs, the abundance and fate of extracellular ARGs in an AnMBR effluent have not been examined. This study elucidates the effect of combined antibiotics loading (ampicillin, erythromycin, and sulfamethoxazole) on the abundance of intracellular and extracellular ARGs in an AnMBR effluent over a period of five weeks. Quantification of targeted genes revealed an overall enrichment of intracellular ARGs (iARGs) and depletion of extracellular (exARGs) in response to antibiotics addition, which suggests exARG uptake as a significant mode of horizontal gene transfer in AnMBR effluents. Comparison of the iARG and exARG abundance profiles showed a potential bias for exARG uptake located on small plasmids compared to large plasmids.

**Importance:** Antibiotic resistance dissemination is facilitated through horizontal gene transfer (HGT) of ARGs. Currently, conjugation is considered to be the dominant mechanism during wastewater treatment. However, recent studies have detected high abundances of exARGs, implying that transformation may play a greater role in dissemination. While previous studies quantified iARGs and exARGs in wastewater treatment facilities, they did not evaluate temporal changes between the two forms. Further, almost no research has differentiated between iARGs and exARGs in anaerobic processes, which are being considered to replace aerobic activated sludge processes. This study specifically investigates the abundance of targeted iARGs and exARGs in AnMBRs in response to antibiotic pressure to quantify potential exchange of ARGs between intracellular and extracellular compartments. Our findings suggest that exARGs located on small plasmids are preferentially taken up by cells under antibiotic pressure compared to large plasmids, which implies heterogenous HGT mechanisms among the plasmid community.

## 1. Introduction

The global spread of antibiotic resistance continues to be a grave threat to human health (Chioro, et al., 2015). Approximately 2 million clinical cases and 23,000 deaths related to antibiotic resistant infections occur annually in the US (Frieden, 2013). Current projections estimate that the yearly worldwide deaths related to antibiotic resistance will reach 10 million by the year 2050 (O’Neil, 2014). While conventional wastewater treatment plants (WWTPs) are effective at managing organics and nutrients, they are not designed or operated to treat emerging contaminants like antibiotic resistance genes (ARGs) and antibiotic resistant bacteria (ARB). With growing awareness of antibiotic resistance, numerous studies have identified WWTPs as hot spots for the propagation of ARGs and ARB (Bouki, et al., 2013, Michael, et al., 2013). Studies on the effluent quality of conventional WWTPs have reported incomplete removal of both ARGs and ARB (McConnell, et al., 2018, Quach-Cu, et al., 2018). Therefore, ARGs and ARB in treated wastewater can continue to proliferate via horizontal gene transfer (HGT) and vertical gene transfer when discharged into receiving environments.

Analyses of sediment microbial communities near outfall sites of WWTPs have detected similar patterns of ARGs and gene classes as those in WWTP effluents (Chu, et al., 2018). While the influence of effluent ARGs on the sediment microbial community is less prominent with increasing distance from effluent outfall sites, the practice of water reuse (e.g., for agricultural irrigation), removes the environmental buffers associated with conventional handling of treated wastewater and results in direct ARGs and ARB loading to receiving soils. Observations of recent studies generally suggest a positive association between proliferation of soil ARGs and contact time with treated wastewater (Fahrenfeld, et al., 2013, Han, et al., 2016, Corno, et al., 2019). Therefore, reducing the abundance of antibiotic resistance elements in treated wastewater is critical to limiting the spread of antibiotic resistance in water reuse applications (Pruden, et al., 2013).

Anaerobic membrane bioreactors (AnMBRs) are an emerging wastewater treatment technology that couples anaerobic biological treatment with membrane separation to recover energy, reduce sludge production, and produce a high-quality effluent comparable to activated sludge processes (Smith, et al., 2012). Although low in carbon, AnMBR effluents are rich in nutrients and can be utilized effectively in agricultural reuse scenarios to offset artificial fertilizer needs. Further, the combination of membrane separation with slower microbial growth under anaerobic conditions can theoretically reduce the load of ARGs and ARB in treated wastewater (Le, et al., 2018). However, the fate of ARGs and ARB within AnMBRs remains poorly understood. Previously, we reported the first long-term investigation of eight ARGs, spanning multiple ARG subtypes, in a bench-scale AnMBR under step-wise increases in influent antibiotic concentrations (Zarei-Baygi, et al., 2019). Our results revealed stark differences in the ARG abundance profile of the biomass versus effluent under all conditions tested. The steep reduction in the ARG abundance profile from the biomass to the effluent is likely due to the effective microbial retention by the submerged membranes(Li, et al., 2019). However, size exclusion by membrane pores may not retain extracellular ARGs (exARGs). Moreover, a growing number of studies have reported a high abundance of extracellular DNA (exDNA) in WWTPs and have suggested that the transformation of exARGs might play a larger role in antibiotic resistance dissemination than previously considered (Mao, et al., 2014, Dong, et al., 2019). Therefore, we suspect that the ARG profile in AnMBR effluent may be heavily influenced by exARGs passing through the membrane pores.

Of concern to human health are the many human pathogens known to enter natural competency (e.g., *Campylobacter*, *Haemophilus*, *Helicobacter*, *Neisseria*, *Pseudomonas*, *Staphylococcus*, *Streptococcus*, *Vibrio*), with some pathogenic strains having a subset of their clonal population in a state of constitutive competency (e.g., *Neisseria gonorrhea* and *Helicobacter pylori).* While the transformation of exDNA may be limited to smaller plasmids and not be effective at disseminating large multi-drug resistant plasmids, smaller non-conjugative plasmids can also confer multi-drug resistance in bacteria (San Millan, et al., 2009, Lean and Yeo, 2017). Therefore, the direct loading of exARGs, iARGs, and ARB to areas of potential human contact could significantly accelerate the development of antibiotic resistant human pathogens.

Previous studies on exARGs and iARGs have focused on investigating their abundance at multiple environmental locations, such as various types of WWTPs, water sources, and sediments (Mao, et al., 2014, Wang, et al., 2016, Zhang, et al., 2018, Dong, et al., 2019). However, no studies to date have examined the effect of antibiotics addition on exARG and iARG abundance over time. The dynamic relationship between exARGs and iARGs within a microbial community in antibiotic-influenced environments remains an area of much needed investigation. The potential cyclical movement of exDNA between microbial hosts and the extracellular compartment may be a fundamental mechanism for the conservation and propagation of ARGs and ARG associated MGEs between distinct and distant microbial communities. While AnMBRs could theoretically lessen the load of ARGs and ARB to receiving environments compared to conventional treatment process, the potential propagation of exARGs in the effluent stream needs further research. In this study, we characterized the effect of a combined mixture of antibiotics loading on the antibiotic resistance characteristics of an AnMBR effluent stream.

## 2. Materials and Methods

### 2.1 AnMBR Configuration and Monitoring

A bench-scale AnMBR with a working volume of 5 L was operated at 25 °C as previously described (Zarei-Baygi, et al., 2019). Briefly, the AnMBR housed three separate submerged flat-sheet silicon carbide microfiltration membranes (0.1 μm pore size, Cembrane, Denmark), with a total effective membrane area of approximately 0.015 m^2^. Synthetic wastewater was used to reduce the variable background influence of antibiotics, ARB, and ARGs associated with real wastewater. The synthetic wastewater recipe was formulated to represent US domestic wastewater (Supporting Information (SI) Table S1) (Smith, et al., 2013).The AnMBR was seeded with sludge from a mesophilic anaerobic digester at the Joint Water Pollution Control Plant (Carson, CA). AnMBR performance was monitored by measuring water quality parameters including soluble chemical oxygen demand (sCOD), total chemical oxygen demand (tCOD), volatile fatty acids (VFAs), and mixed liquor and volatile suspended solids (MLSS/MLVSS). Biogas production was monitored by an in-line gas flowmeter and assessed for methane content by gas chromatography with flame ionization detection (GC-FID). Steady AnMBR performance was defined as low effluent COD (<50 mg/L), stable biogas production, and high methane content (>60%) over a phase of 10 days. After reaching steady AnMBR performance, three antibiotics, sulfamethoxazole (sulfonamide), erythromycin (macrolide), and ampicillin (β-lactam), were added to the influent at 250 μg/L (day 3) to represent antibiotic concentrations at the high range of hospital wastewater(Xu, et al., 2016, Kulkarni, et al., 2017). Influent antibiotics concentration was maintained at 250 μg/L for the entirety of the antibiotics loading phase (day 3 to day 35). Effluent lines were cleaned with sodium hypochlorite 0.5% (v/v) before the start of the experiment. Membrane modules were removed for physical and chemical cleaning 0.5% (v/v) before the addition of antibiotics.

### 2.2 Antibiotic Quantification

Antibiotic quantification procedures were carried out as previously described (Zarei-Baygi, et al., 2019). Briefly, individual antibiotic concentrations of each sample were analyzed by direct injection liquid chromatography mass spectrometry with electrospray ionization (LC-ESI-MS) on a 6560 Ion Mobility Quadrupole Time-of-Flight (IM-QTOF) LC-MS system (Agilent). Chromatographic separation and ionization were achieved by employing a 1290 Infinity UHPLC with EclipsePlus C18 column (2.1 mm; 50 mm; 1.8um) followed by a Dual Agilent Jet Stream (ASJ) ESI. Standard curves were generated by matrix-matched external calibration of serial dilutions of antibiotics purchased from Sigma (>99% purity). Practical quantitation limits (PQL) for each target compound were determined to be <0.1 μg/L based on previously optimized LC-ESI-MS conditions (Zarei-Baygi, et al., 2019). Details on sample preparation, optimized LC program, and MS operational conditions can be found in the (SI Table 3).

### 2.3 qPCR quantification of ARGs

qPCR reactions were carried out using a LightCycler 96 (Roche, Basel, Switzerland) targeting a set of nine genes. The nine targeted genes included: a class one integrase gene (*intl*1), a single copy per cell gene (*rpo*B), and seven ARGs conferring resistance to β-lactams (*oxa*-1 and *amp*C), macrolides (*erm*F), sulfonamides (*sul1* and *sul2*), and tetracyclines (*tet*W and *tet*O). Targeted ARGs were chosen based on their common detection in domestic wastewater studies (Pruden, et al., 2006, Ma, et al., 2011, Munir, et al., 2011). qPCR reactions were done in 20 μL reactions with 10 μL of qPCR master mix (Forget-Me-Not EvaGreen, Biotium, Fermont, CA), forward and reverse primers at 0.25 μM (final concentration) each, 1 μL of DNA template, and ddiH_2_O. Each reaction was performed in triplicate. Details for the thermal cycling conditions for all targeted ARGs are provided in the (SI Table 2). All qPCR results were normalized to effluent sample volume for comparison of ARG abundance across sampling points. Results were represented as total abundance instead of normalized values relative to chromosomal *rpoB* gene due to previous reports of faster decay rates for chromosomal DNA versus plasmid DNA, which could lead to inflated reports of exARG concentration when normalizing to chromosomal genetic markers (Mao, et al., 2014).

### 2.4 Intracellular and extracellular DNA extraction

iDNA and exDNA extractions were carried out using the same 500 mL effluent samples for each sampling point. Internal standards to correct for DNA recovery across sample processing were added to each effluent sample by spiking with approximately 2×10^6^ copies of plasmid pUC19 immediately after collection. Spiked effluent samples were filtered using vacuum filtration through sterile cellulose acetate membrane filters (0.22um, 45mm diameter, Whatman). Processed filters with collected biomass were used for iDNA extractions. The processed filters were cut into pieces and placed in 2 mL sterile tubes and mixed with lysis buffer before bead beating with zirconium beads. iDNA extractions were carried out using the Maxwell 16 instrument (Promega) and eluted in 100μl of buffer AE. The filtrates were used for exDNA extractions using nucleic acid adsorption particles (NAAP) adapted from Wang et al (Wang, et al., 2016). Briefly, autoclaved Al(OH)_3_ solutions (47.8%, V/V) were mixed with 5% (g/mL) silica gel (60-100 mesh size, Sigma) and dried for 36 hours at 60°C. Dried silica gels coated with Al(OH)_3_ were then sealed in a cylindrical glass container (1.5 x 40 cm) as NAAP columns. The previously mentioned filtrate samples were passed through the NAAP columns. Adsorbed exDNA were eluted from the NAAP columns with 100 mL of elution buffer (15 g/L NaCl, 30 g/L tryptone, 15 g/L beef extract, 3.75 g/L glycine, 0.28 g/L NaOH, pH of 9.3 ± 0.2). Eluates were then collected and filtered with polyethersulfone filters (0.22 um, Millipore, USA). ExDNA in the filtrates were precipitated using an equal volume of isopropanol, incubated at room temperature for 16 h, and centrifuged at 10,000g for 10 min at room temperature. After decantation of the supernatant, the centrifuged pellets were mixed with 70% ethanol (v/v) and centrifuged again at 10,000 g for 5 min at room temperature. After a second decantation, the residual ethanol was evaporated in a 60 °C oven and the pellets were re-suspended in 4 mL of sterilized TE buffer. Extracted DNA samples were quantified using Quant-iT PicoGreen (Thermo Fisher) and a BioSpectrometer (Eppendorf, Hamburg, Germany). iDNA extraction efficiency was carried out without the use of an iDNA internal standard due to the consistent performance of the Maxwell 16 instrument (Promega) (data not shown). Extraction efficiency of each exDNA extraction was assessed using spiked pUC19 serving as the exDNA standard (SI Table 2). The qPCR results for each exARG data point were adjusted based on the corresponding extraction efficiency of the exDNA standard (e.g,, gene copies of exampC/(gene copies of pUC19/total spiked gene copies of pUC19)). All DNA extracts were stored at −80 °C until analysis.

### 2.5 Statistical Analysis Methods

Statistical analyses were performed using MaxStat Lite 3.6. Significant changes in ARG abundance at different time points were assessed using a 2-tailed unpaired student’s t-test. Pearson and Spearman rank correlations were used to assess correlations between data points over a 95% confidence interval. Strength of correlations were identified based on the Pearson coefficient r, as r > 0.7 or r < −0.7 for strong correlations, −0.7 <r<−0.5 or 0.5<r<0.7 for moderate correlations, and −0.5 < r < −0.3 or 0.3 < r < 0.5 for weak correlations. Pearson correlations were performed on separated iARG and exARG data sets. Spearman rank correlations were performed on the combined iARG and exARG data sets due to the dynamic nature of gene transport, synthesis, and degradation.

### 2.6 Heterotrophic Plate Counts

Total bacteria and ARB in the effluent were enumerated using the heterotrophic plate count method with nutrient agar as the media (Federation, 2005). Ampicillin (AMP), tetracycline (TET), erythromycin (ERY), and sulfamethoxazole (SMX) resistant bacteria counts were determined by growth on nutrient agar plates supplemented with corresponding AMP, TET, SMX, or ERY. TET resistance bacteria counts were included as a control given the lack of tetracycline in the synthetic wastewater. Final concentration for each antibiotic was guided by MIC reports in previous studies: AMP, 16 μg/mL; TET, 16 μg/mL; ERY, 50.4 μg/mL; and SMX, 18.1 μg/mL(Pei, et al., 2006, Zheng, et al., 2017). All plates were incubated at 35°C for 48 hrs.

## 3. Results and Discussion

### 3.1 Stable Reactor Performance Under Mixed Antibiotics Loading

Effluent COD averaged 45.4 ± 10.7 mg/L, equating to a COD removal of 90.0 ± 1.8% throughout the experimental phase. Average biogas production and methane content were 736 ± 20 mL/d and 536 ± 14 mL/d, respectively. The average MLSS and MLVSS were 10.6 g/L ± 1.3 and 9.6 ± 0.3 g/L, respectively. The addition of mixed antibiotics (SMX, ERY, and AMP), at 250 ug/L each, to the influent showed minimal perturbation to the reactor performance (SI Fig1), except for day 17 which saw a slight decrease in biogas production. These results are consistent with previous studies which have shown that this range of antibiotic loading on anaerobic reactors is well below the threshold that would cause a disruption in system performance (Aydin, et al., 2015, Xiong, et al., 2017). Antibiotic removal was relatively stable across the operational phase, with AMP removal ranging from 89-98%, followed by SMX at 69-78%, and ERY at 40-58% (Fig 1). The wide-ranging specific removal rates of these antibiotics in the AnMBR are consistent with those observed for other mainstream anaerobic wastewater treatment systems, which can vary greatly in comparison to conventional aerobic treatment schemes (Harb, et al., 2019).

**Figure 1:**
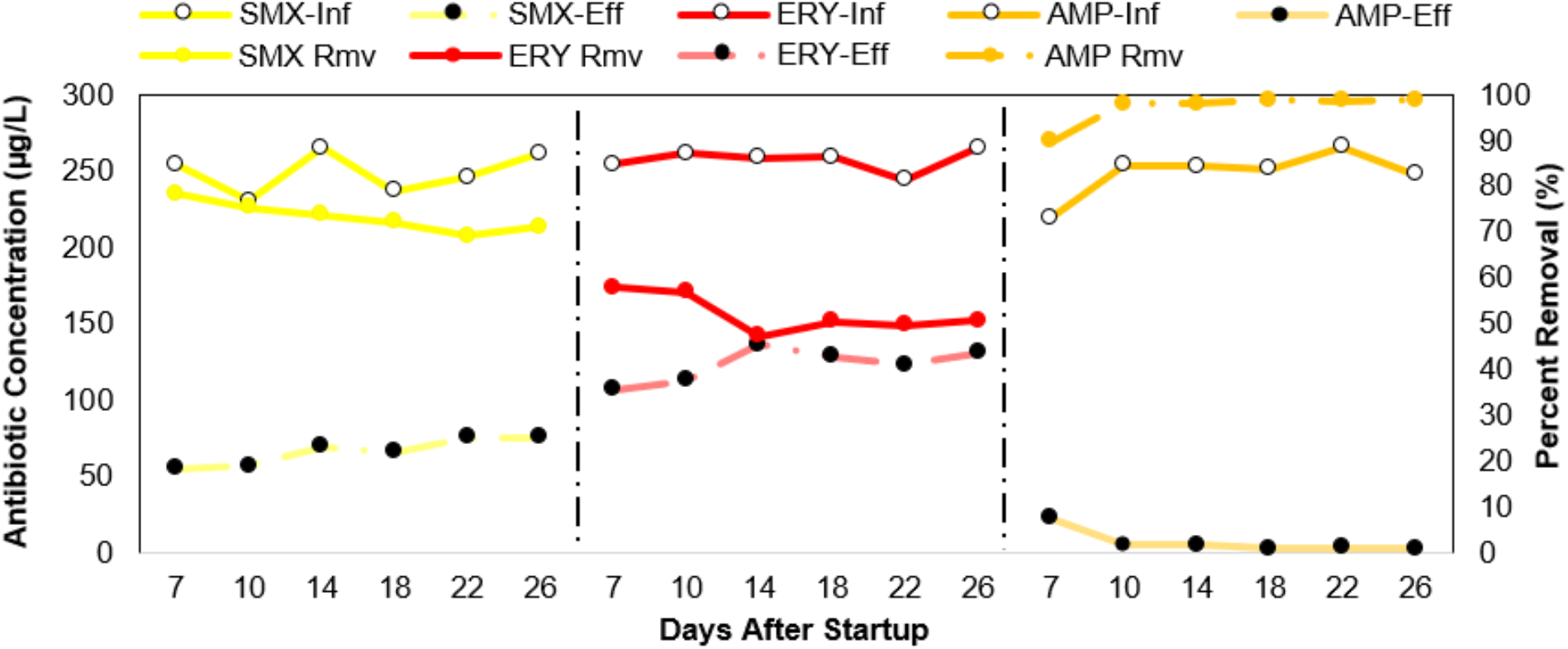
Measured antibiotic concentrations in the influent and effluent of AnMBR along with antibiotic removal percentage. Antibiotics (ampicillin, erythromycin, and sulfamethoxazole) were added to influent at a final concentration of 250 µg/L from day 3 to day 35.

### 3.2 SMX Resistant Bacteria Count Approaches Total Culturable Bacteria Count

Overall, SMX resistance showed the greatest degree of proliferation among the types of antibiotic resistance tested. SMX resistant bacteria in the effluent increased by > 3 log (CFU/mL) (p=0.032) over the course of the experiment, with the number of total bacteria increasing by 0.68 log (CFU/mL) (p=0.027) over the same duration (Fig 2). The ratio of SMX resistant bacteria to total bacteria exhibited a non-linear rise from 55.5% (day 1) to 96.5% (day 57). In comparison, the ratio of AMP resistant bacteria to total bacteria rose from 68.7% to 75.4% and the ratio of ERY resistant bacteria to total bacteria fell from 89.3% to 76.7%. As expected, (given no addition of TET in the influent) counts for TET resistant bacteria frequently fell below the limit of detection (LOD= 10 CFU/mL). Among the ARB types tested, only SMX resistant bacteria showed a positive association between the increase in effluent abundance and antibiotic dosing. Our findings were consistent with our previous report and Le et al., where sulfamethoxazole was found to be one of three antibiotics to increase with its corresponding ARB among 19 targeted antibiotics in an MBR system (Le, et al., 2018, Zarei-Baygi, et al., 2019). It is important to note that heterotrophic plate counts only capture a small fraction of microbial diversity. Further, aerobic plating of effluents from anaerobic systems favors quantification of facultative microorganisms and excludes detection of obligate anaerobes. Despite these methodological limitations, HPC used in this capacity still provides useful data regarding relative changes in ARB, which complements our parallel use of culture-independent methods (i.e., qPCR).

**Figure 2:**
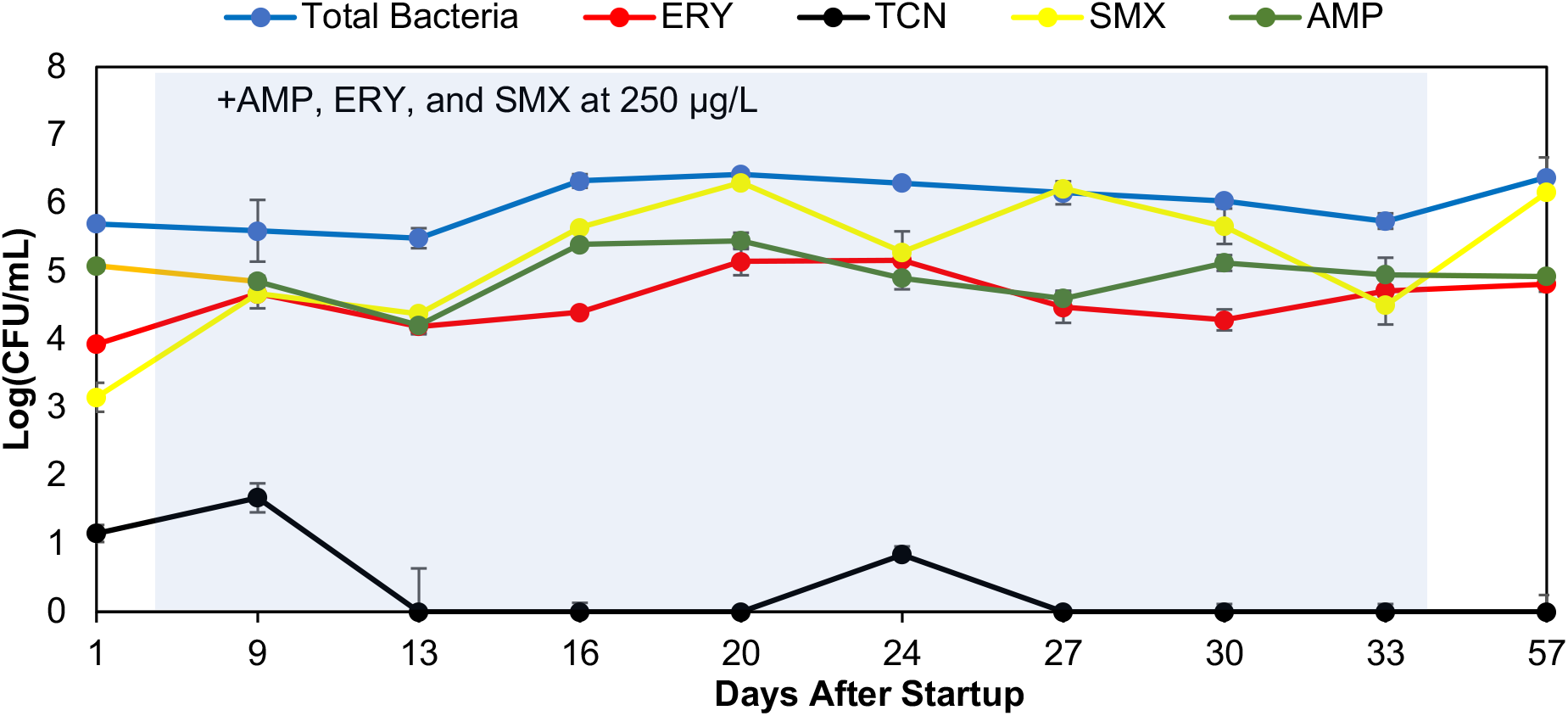
Heterotrophic plate count for effluent total bacteria, erythromycin resistant bacteria (ERY), tetracycline resistance bacteria (TET), sulfamethoxazole resistant bacteria (SMX), and ampicillin resistant bacteria (AMP) over the experimental run. All samples were plated in duplicate. The addition of antibiotics lasted from day 3 to day 35.

### 3.3 Total iARGs and exARGs Showed Varied Response Towards Antibiotics Addition

Total iARGs were consistently more abundant than total exARGs by at least 2 log (Gene Copies/mL) throughout the experimental period (Fig 3). Similar findings of higher iARG versus exARG abundance have been reported in most aquatic environments examined (Nielsen, et al., 2007, Mao, et al., 2014, Zhang, et al., 2018, Hao, et al., 2019). In contrast, studies on sediment samples frequently report higher abundance of exARGs than iARGs (Mao, et al., 2014, Dong, et al., 2019). The higher abundance of exARGs versus iARGs in sediment samples is likely due to the adsorption of exDNA to soil colloids, clay particles, and organic matter that can decrease the susceptibility of exDNA degradation from nuclease attacks (Crecchio and Stotzky, 1998, Demanèche, et al., 2001).

**Figure 3:**
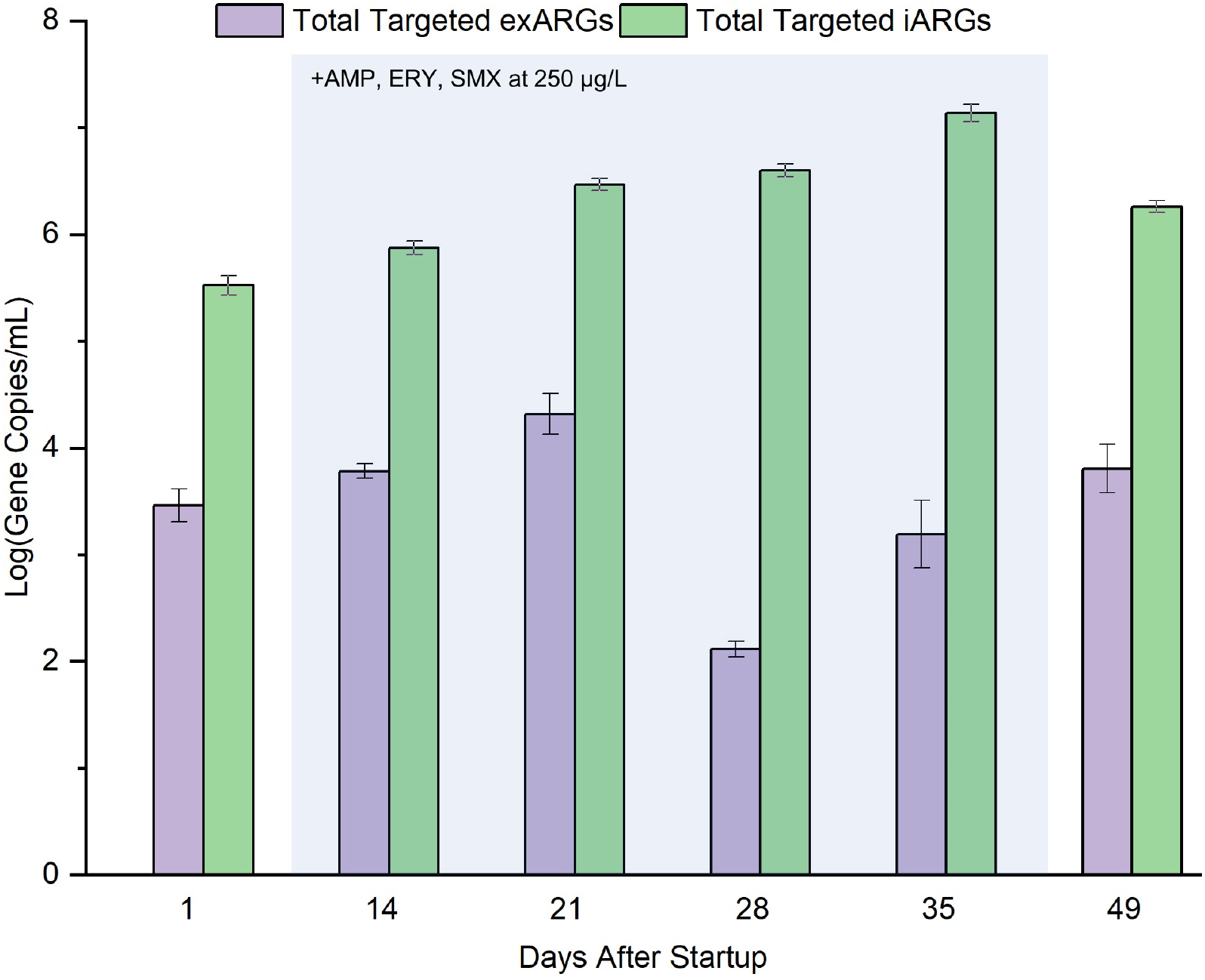
Abundance of total targeted intracellular and extracellular ARGs over the course of the experiment. Targeted ARGs consisting of *amp*C, *erm*F, *sul*1*, sul*2, *tet*W, *tet*O, and *oxa-*1 were summed for each sampling date and plotted as log(Gene Copies/mL). The mixed antibiotics period lasted from day 3 to day 35.

The addition of antibiotics in the influent coincided with increasing abundance of total iARGs in AnMBR effluent, rising from 5.52 (day 1) to 7.13 (day 35) log (Gene Copies/ml). The observed increase in total iARGs during the antibiotics loading phase is likely due to the selection for ARB and their associated ARGs in the effluent stream. The abundance of total exARGs, over the same time frame, exhibited a non-linear decrease from 3.46 (day 1) to 3.16 (day 35). Interestingly, total exARGs showed an initial rise to 4.2 log (gene copeies/mL) (day 21) followed by a precipitous drop to 2.11 log (Gene Copies/ml) midway through the experimental run (day 28). The initial increase in total exARGs before day 28 could be due to antibiotic induced exDNA release for the purpose of biofilm formation and HGT (Zafra, et al., 2012, Okshevsky and Meyer, 2015, Sugimoto, et al., 2018). Moreover, antibiotics, biofilm formation, and exDNA are all associated with the induction of natural competency of many bacterial species (Li, et al., 2001, Ibanez de Aldecoa, et al., 2017). Therefore, an increase in competent microbial members within the effluent stream could explain the sharp decrease in total exARGs abundance on day 28. In the post-antibiotics loading phase, the increase in total exARGs is likely due to the loss of ARG-associated MGEs as the fitness advantage of retaining ARG-associated MGEs decreases. However, the persistence of MGEs in microbial hosts under antibiotic-free conditions is influenced by the surrounding genes. While low copy large plasmids typically encode for additional maintenance genes (e.g., partitioning and toxin-antitoxin systems) that promote their own survival in the absence of antibiotic pressure (Ghaly and Gillings, 2018), high copy small plasmids do not encode for these maintenance genes and are generally more easily lost in the absence of strong selective pressure.

### 3.4 Mixed-Antibiotics Loading Increases the Abundance of Effluent iARGs

In the pre-antibiotics phase, *intl*1 was the most abundant target gene detected in the effluent, followed by *sul*1, *rpoB*, *sul*2, *tet*O, *tet*W, *erm*F, *oxa*-1, and *amp*C (Fig 4). During the antibiotics loading phase, the abundance of *sul*2 exhibited the highest increase, rising from 4.57 to 7.08 log (Gene Copies/ml) (p<0.01). The abundance of s*ul*1 and *intl*1 increased from 5.9 to 6.56 log (Gene Copies/ml) (p=0.168) and from 5.46 to 6.21 log (Gene Copies/ml) (p=0.238), respectively. The relatively high abundance of *sul*1, *sul*2, and *intl*1 among targeted ARGs in our study is consistent with previous reports (Rowe, et al., 2016, Xu, et al., 2016, Zarei-Baygi, et al., 2019). Interestingly, the abundance of *oxa-*1 rose from 2.39 to 3.58 log (Gene Copies/ml) (p < 0.01), while the abundance of *amp*C decreased from 1.16 to 0.9 log (Gene Copies/ml) (p=0.02). The preferential selection of *oxa*-1 over *amp*C could be due to the co-localization of *oxa*-1 and *sul*2 on the same MGE, which would allow for the co-selection of *oxa-*1 from the SMX addition. This hypothesis is supported by the strong correlation between *sul*2 and *oxa*-1 (r= 0.772) (p<0.01) and the low AMP (2.8 µg/L to 22.45 µg/L) selective pressure in the effluent. Despite high ERY concentrations (107.0 µg/L to 130.5 µg/L) in the effluent, *erm*F abundance showed modest increase from 2.91 to 3.86 log (Gene Copies/ml) (p<0.01).

**Figure 4:**
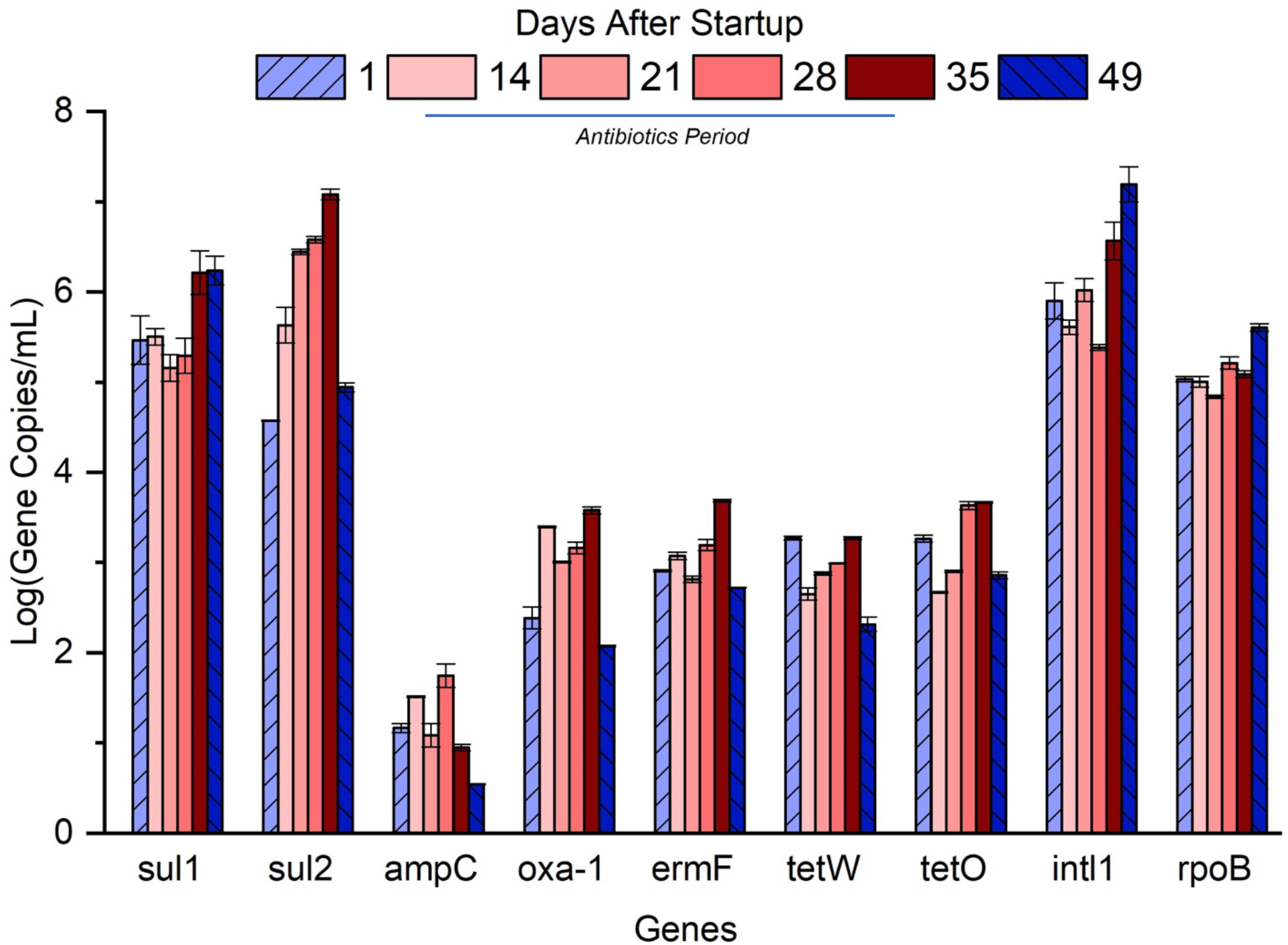
Abundance of targeted intracellular ARGs, *int*I1, and *rpo*B (Gene Copies/mL) in the AnMBR effluent. Error bars represent the mean value and standard deviations of triplicate qPCR runs for each sample. Antibiotics (ampicillin, erythromycin, and sulfamethoxazole) were added to influent at a final concentration of 250 µg/L from day 3 to day 35.

In the post-antibiotics phase, *sul*2 levels significantly dropped by 2.16 log (Gene Copies/ml) (p<0.01). Consistent with our previous co-localization hypothesis, the abundance of *oxa*-1 showed a similar sharp decrease, dropping from 3.5 to 2.08 log (Gene Copies/ml) (p<0.01). Additional iARG decreases included *erm*F, *tet*O, and *tet*W, which declined in abundance from 3.68 to 2.72 log (Gene Copies/ml) (p <0.01), 3.27 to 2.31 log (Gene Copies/ml) (p <0.01), and 3.67 to 2.86 log (Gene Copies/ml) (p <0.01), respectively. Surprisingly, the abundance of *sul*1 and *intl*1 remained elevated at 6.59 and 7.19 log (Gene Copies/ml), respectively. The elevated concentrations of *sul*1 and *intl*1 genes in the post antibiotics phase may be due to the greater persistence of low copy number large plasmids compared to high copy small plasmids in microbial hosts (Carroll and Wong, 2018). On a population level, non-conjugative plasmids tend to be smaller in size and are present in higher copy numbers (up to 200 copies per cell) compared to conjugative plasmids, which tend to be much larger in size and present in lower copy numbers (<10 copies per cell) (Watve, et al., 2010, Shintani, et al., 2015). Under selective pressure, high copy number plasmids could theoretically increase in abundance within a microbial community more rapidly than low copy plasmids. However, small non-conjugative plasmids typically do not encode for maintenance or stability genes and are easily lost when the selective pressure is removed. In contrast, low copy conjugative plasmids may exhibit longer persistence within a microbial community due to additional maintenance systems encoded on the plasmid backbone (e.g., partitioning and toxi-antitoxin systems) (Garcillan-Barcia, et al., 2011). Overall, *sul*1 and *sul*2 were the two most abundant ARGs detected and *sul*2 exhibited the greatest increase under antibiotic pressure. Further, the sharp rise of *sul*2 in response to antibiotic stress and the persistence *of sul*1 in the post-antibiotics phase is consistent with the SMX resistant bacteria proliferation observed in our HPC results.

Pearson correlation analysis of the iARG data set revealed significant correlations between two sets of genes, *sul*2, *oxa*-1, and *tet*O (r>0.65, p<0.05) and *erm*F, *sul*1, and *intl*1 (r > 0.6, p <0.05), which suggests the presence of multi-drug resistant plasmids and/or distinct modes of plasmid propagation rates (e.g., low versus high copy plasmid). Moreover, *sul*1 was strongly correlated with *intl1* (r = 0.869, p < 0.01), while no correlation was found between *sul*2 and *intl*1 (r = −0.059, p =0.853), which suggests that the discrepant response of *sul*1 and *sul*2 to the addition of antibiotics was attributed to the different types of MGEs surrounding *sul*1 and *sul*2. Consistent with this explanation, previous studies examining the characteristics of isolated plasmids harboring *sul*1 and *sul*2 commonly detected *sul*2 on small non-conjugative plasmids whereas *sul*1 was exclusively found on large conjugative plasmids (Enne, et al., 2004, Antunes, et al., 2005, San Millan, et al., 2009, Wu, et al., 2010, Dominguez, et al., 2019). The strong correlation between *erm*F with *sul*1 and *intl*1 suggest that *erm*F is either co-localized with *sul*1 and *intl*1 or located on similar low copy number conjugative plasmids, which could explain the modest increase in abundance for *sul*1, *int*1, and *erm*F despite significant concentrations of SMX (54.9 µg/L to 75.2 µg/L) and ERY (107.0 µg/L to 130.5 µg/L) in the effluent stream.

### 3.5 Temporal Abundance Prolife of Effluent exARGs and iARGs Revealed Distinct Patterns

In the pre-antibiotics phase, all exARGs were detected, ranging from 0.25 to 3.1 log (Gene Copies/ml) (Fig 5). Interestingly, the addition of antibiotics coincided with an overall decrease in exARG abundance, which contrasted the enrichment of iARGs during the same period. Similar to iARGs, exARGs are likely associated with MGEs of varied sizes. The uptake of exDNA by microbial hosts has been documented to favor small plasmids and DNA fragments, which could cause the overall variability of exARG abundance (Prudhomme, et al., 2006, Slager, et al., 2014). Therefore, the size of the genetic carrier harboring exARG would influence its rate of degradation and uptake by microbial hosts.

**Figure 5:**
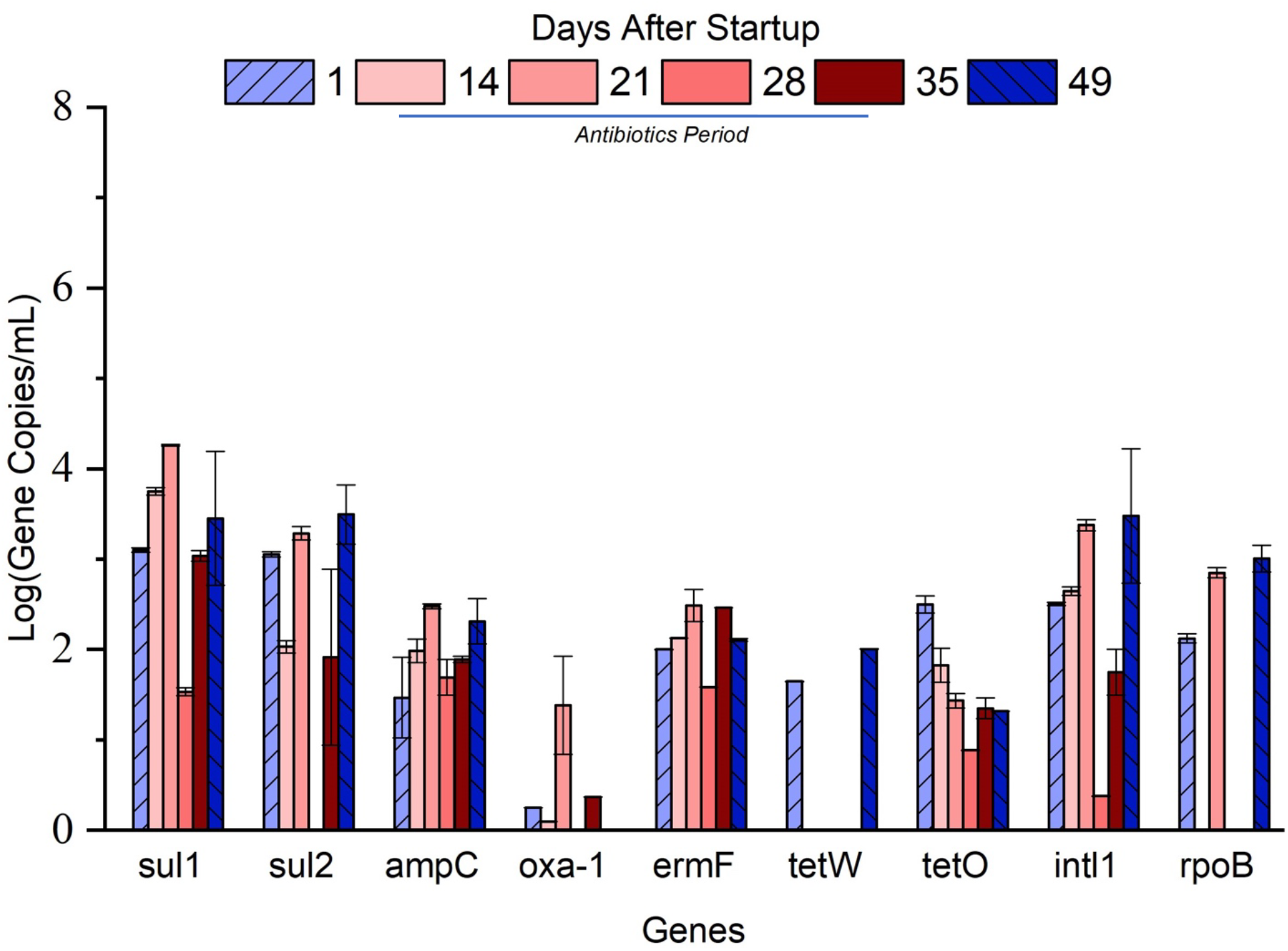
Abundance of targeted extracellular ARGs, *intl*1, and *rpo*B log(Gene Copies/mL) in the AnMBR effluent during the course of the experimental run. Error bars represent the mean value and standard deviations of triplicate qPCR runs for each sample. Antibiotics (ampicillin, erythromycin, and sulfamethoxazole) were added to influent at a final concentration of 250 µg/L from day 3 to day 35. Detection limit for each ARG are provided in (SI Table 2)

Pearson correlation analysis of the exARG data set revealed distinct patterns of removal, which supports the hypothesis of size dependent exDNA uptake. Similar to the Pearson correlation analysis for iARGs, extracellular *sul*1 was strongly correlated with extracellular *intl*1 (r= 0.87, p<0.01). Interestingly, extracellular *sul*2 was strongly correlated with extracellular *intl*1 as well (r=0.78 p<0.01) and was moderately correlated with extracellular *sul*1 (r=.59 p<0.01). While some studies have identified *sul*2 to be distinctly separate from *sul*1 and *intl*1, a few reports have detected *sul*2 genes on large conjugative plasmids along with *sul*1 and *intl*1 genes (Phuong Hoa, et al., 2008, Wu, et al., 2010, Wu, et al., 2010). The positive correlation of extracellular *sul*1, *sul*2, and *intl*1 in the extracellular compartment could be due to the lower uptake efficiency of large plasmids compared to small plasmids. Since *sul*2 has been documented on both large and small plasmids, there could be a preferential uptake of small *sul*2-associated plasmids while leaving behind larger *sul*2-associated plasmids. From the iARG profile, intracellular *erm*F, *sul*1, and *intl*1 all exhibited propagation patterns resembling those of low copy number conjugative plasmids, which are generally larger than non-conjugative plasmids (Garcillan-Barcia, et al., 2011). Since exARGs are simply the cell-free form of iARGs, the greater persistence of extracellular *erm*F, *sul*1 and *intl*1 over extracellular *sul*2, *oxa*-1, and *tet*O could be due to the lower uptake efficiency of larger plasmids compared to smaller plasmids.

Spearman rank correlation analysis was performed on the combined iARG and exARG data sets to further asses the temporal relationship between iARGs and exARGs. Among the ARGs examined, only extracellular *sul*2 and extracellular *tet*W were negatively correlated to intracellular *sul*2 (r=−0.68, p<0.01) and intracellular *tet*W (r = −0.836, p <0.01), respectively. The negative correlation between extracellular *sul*2 and intracellular *sul*2 is consistent with our hypothesis that exARGs can be taken up and enriched within microbial hosts. The lack of a significant correlation among the remaining exARGs and their intracellular counterpart could be due to additional barriers involved in acquiring and maintaining exogenous genes. As stated previously, the uptake of exDNA by microbial hosts would likely favor small plasmids and DNA fragments Further, the maintenance of exogenous genes, including ARGs, on linear DNA fragments would require recirculation or integration into the microbial host genome through homologous recombination or transposable elements. In general, non-integrated or circularized fragments would likely be degraded for metabolic purposes or released back into the environment. Therefore, the uptake of exDNA could be the first step in acquiring new functional traits and the quality and compatibility of the genetic carrier adds further selection to the maintenance of the acquired genes. Our findings add support to the hypothesis that the extracellular compartment within microbial communities can serve as a reservoir for genetic resources, including ARGs, with potential uptake biases toward small plasmids or DNA fragments.

While *in vitro* studies attempting to induce natural competency for many bacterial species have shown limited success, the complexities of environmental conditions are difficult to recreate in controlled laboratory conditions. Further, the induction of natural competency for some bacterial strains have been shown to rely on the production of signaling molecules from other bacterial species (Zhu, et al., 2011). In the context of wastewater reuse, treated wastewater could provide high concentrations of exARGs, environmental stressors, and signaling molecules from highly complex microbial communities that could promote a feedback loop for exDNA release, biofilm formation, and natural competency in the receiving microbial community. This cyclical uptake and release of exARGs, particularly in plasmid form, could increase the total abundance of exARGs within the extracellular matrix due to multiple replication cycles of plasmids within microbial hosts. Additional studies are needed to unravel the complex dynamics between exARG uptake and release within receiving environments such as soil microbial communities.

In this study, we have shown that under mixed-antibiotics loading (SMX, ERY, and AMP at 250 µg/L), SMX resistant bacteria increase from 55.5% to 96.5% of total culturable bacteria in the effluent. Complimentary qPCR analysis of iARG abundance revealed a stark increase in *sul*2 and the persistence of *sul*1 abundance, which is consistent with our observation of the dominance of SMX resistant bacteria in HPC results. Finally, qPCR analysis of the exARG abundance over the same duration revealed an inverse trend for most exARGs and their intracellular counterparts. Most notably, extracellular *sul*2 was negatively correlated to intracellular *sul*2 (r= −0.68, p=0.001), which suggests that competent cells can acquire antibiotic resistance from exARGs. However, it is important to note that qPCR assays are limited to the quantification of pre-determined genes. Further, the correlations drawn in this paper would benefit from experiments directly tracking the movements of iARGs and exARGs. Future studies should use metagenomic guided qPCR analysis to analyze ARGs pertinent to their research question(s) along with their associated MGEs.

## Supporting Information

**SI Figure 1:**
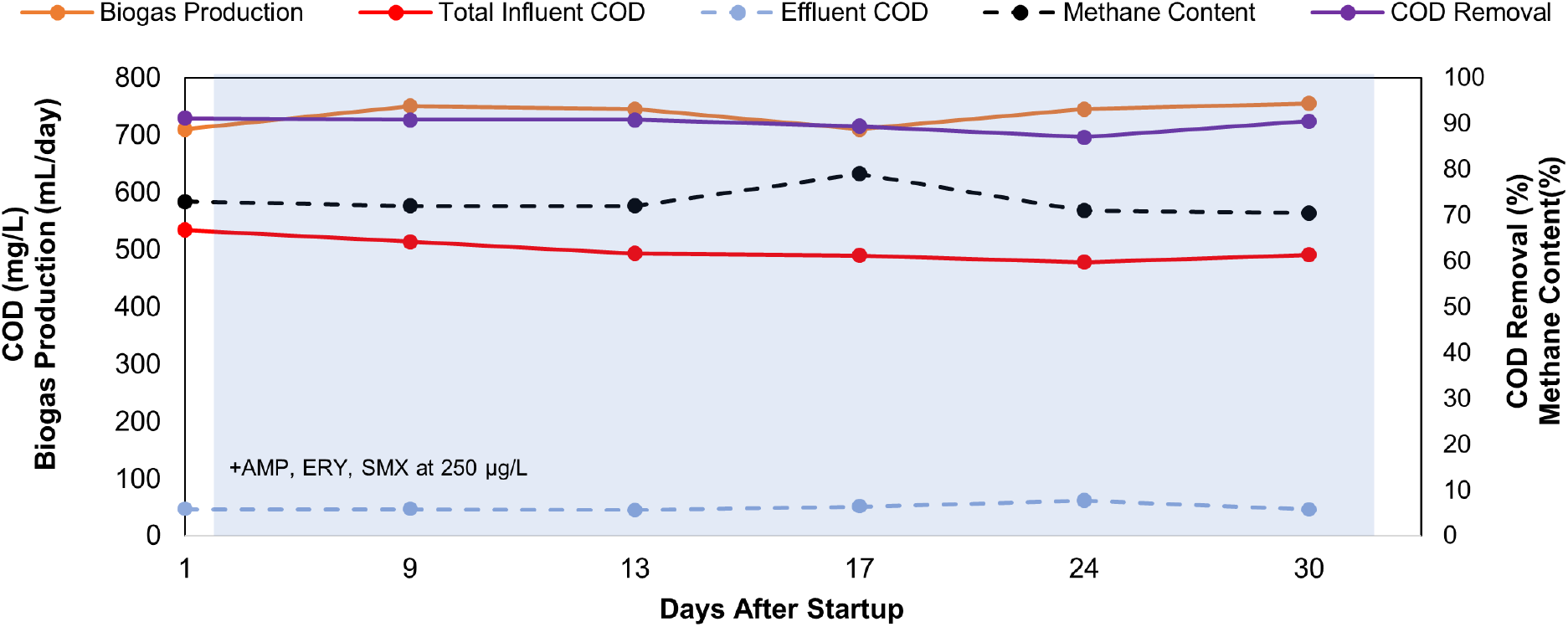
AnMBR performance for COD removal and biogas production during the addition of antibiotics. Biogas production, total COD and effluent COD concentrations are plotted on the primary y-axis. COD removal and methane content are plotted on the secondary y-axis. Antibiotics (ampicillin, erythromycin, and sulfamethoxazole) were added to influent at a final concentration of 250 µg/L from day 3 to day 35.

### Bench-scale anaerobic membrane bioreactor operation

The bench-scale AnMBR contained 5 liters of sludge with a MLSS and MLVSS 7.9±0.4 g/L and 6.1 ± 0.5 g/L, respectively. The influent consisted of synthetic wastewater, which was prepared twice a week. The synthetic wastewater contains two components. A concentration solution and a dilution water. The concentrate solution and dilution water were combined at a ratio of 1:9 to achieve the final concentration for the feed. Table S1 lists the components and final concentration for both solutions.

**SI Table S1:** Synthetic wastewater composition

| Concentrate solution |  | Dilution water |  |
| --- | --- | --- | --- |
| Composition | Concentration (mg/L) | Composition | Concentration (mg/L) |
| Ammonium Chloride | 11.5 | Sodium Bicarbonate | 369 |
| Calcium Chloride | 11.5 | Magnesium Phosphate | 30.8 |
| Iron Sulfate | 7.7 | Potassium Phosphate | 13.8 |
| Sodium Sulfate | 11.5 | Sodium hydroxide | 18.5 |
| Sodium Acetate | 27 |  |  |
| Urea | 87 |  |  |
| Peptone | 11.5 |  |  |
| Yeast | 46 |  |  |
| Milk Powder | 115.4 |  |  |
| Soy Oil | 13.5 |  |  |
| Hydrochloric acid | 0.2 |  |  |
| Starch | 115.4 |  |  |
| Chromium Nitrate | 3.7 |  |  |
| Copper Chloride | 2.5 |  |  |
| Manganese Sulfate | 4.9 |  |  |
| Nickel Sulfate | 1.2 |  |  |
| Lead Chloride | 0.5 |  |  |
| Zinc Chloride | 1.2 |  |  |

Membrane modules were physically cleaned and submerged in 0.5% (v/v) NaOCl solution overnight. In addition, a peristaltic pump was used to continuously circulate the 0.5% NaOCl solution through the membranes. To return the permeate back to neutral pH, the membrane modules were pumped with DI water for 3 hours. During the DI water pumping period, the permeate flux and transmembrane pressure (TMP) results did not indicate any signs of irreversible fouling throughout the experimental run.

A flowmeter (GFM17 Flow Meter, Aalborg, Orangeburg, NY) was used to measure the production of total biogas. The recirculation of headspace biogas through the sparging tubes underneath the membrane modules (rate = 30 ml/min) served to scour the surface of the membranes and prevent membrane fouling. The working volume of the reactor was maintained using an automatic level float switch. Effluent permeate flow was controlled at a rate of 8 min filtration and 2 min backwashing per 10 min period using a peristaltic pump (BT100-1L Multi-channel Peristaltic Pump, Longer, China). Pressure transducers measured the TMP of each membrane module. The permeate flux was set at 7 LMH, which led to hydraulic retention time (HRT) of 16 hours. No sludge was wasted except for sampling, which resulted in a solids retention time (SRT) of 300 days. The software LabVIEW 2014 (Student Edition) was used to monitor and record all relevant AnMBR operational parameters.

Chemical oxygen demand (COD) was measured in accordance with USEPA Method 410.4 using a HI801 Spectrophotometer (Hanna Instruments, Woonsocket, RI, USA). Volatile fatty acids (acetate, propionate, formate and valerate), sulfate, phosphate and chloride were measured by ion chromatography on an ICS 2100 (Thermo Fisher Scientific, Waltham, MA) using methods described previously (Chen and Smith, 2018). Headspace biogas samples and effluent were analyzed using a Trace 1310 GC system (Thermo Scientific, NY) with flame ionization detection (FID) as described previously (Chen and Smith, 2018).

**SI Table S2:**
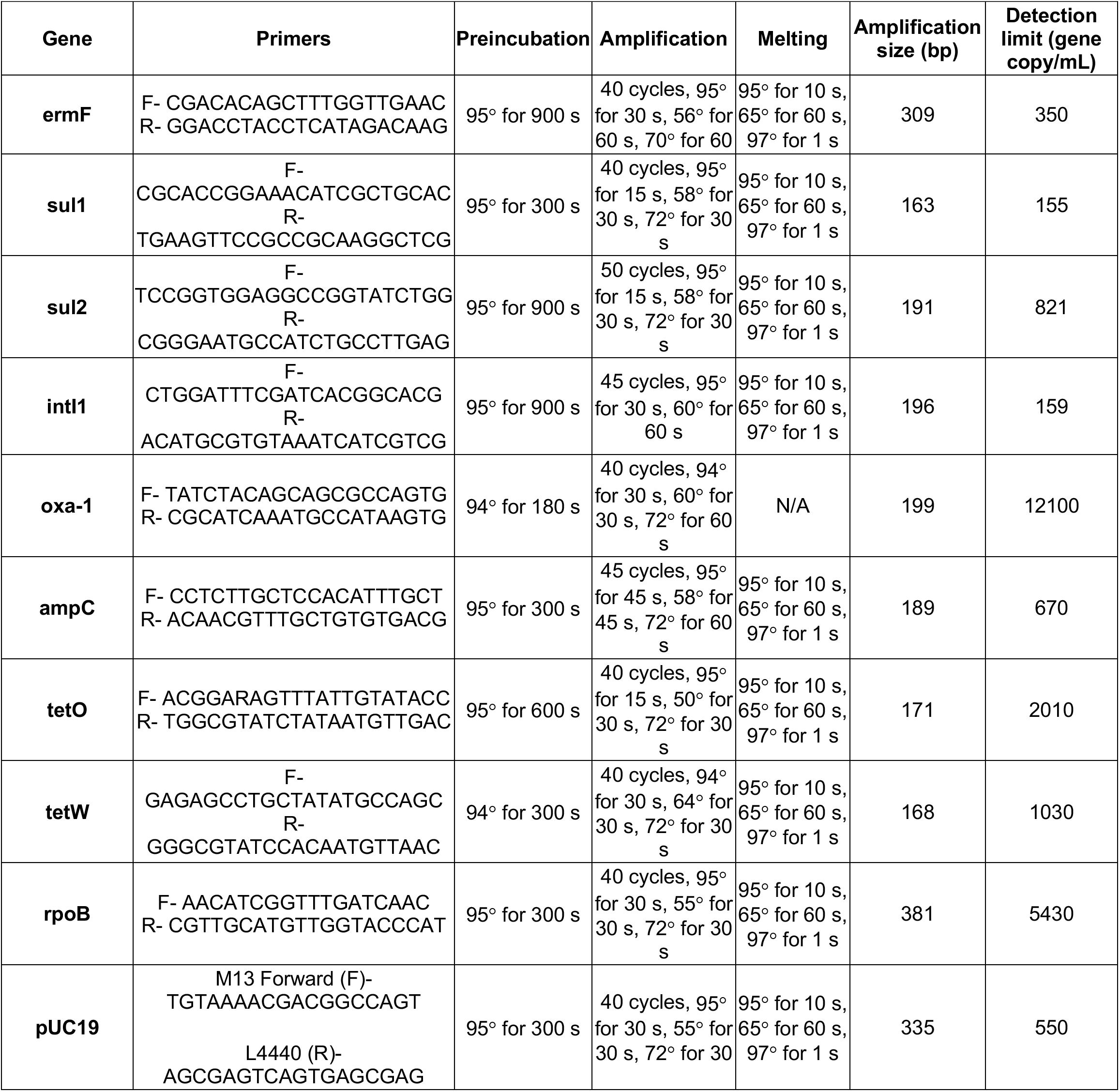
qPCR thermocycling conditions for all forward and reverse primers used in this study

### Antibiotic quantification

All glassware used for antibiotic quantification were baked at 400 ºC for a at least one hour and washed with methanol prior to use. Influent and effluent samples (10 mL) and standard solutions were filtered using a 0.2 µm PTFE syringe filters (Whatman) and a 10 mL syringe with Luer lock tip. Prepped samples were preserved in certified 2 mL amber LC vials (Agilent) and stored at 4 ºC for no more than 3 days prior to analysis. Stock solutions of SMX and ERY were prepared in HPLC-grade methanol at concentrations of 20 mg/L and stored at −20 ºC. AMP stock solution was prepared in HPLC-grade water at 20 mg/L because of its insolubility in methanol and stored at 4 ºC. Five-point standard calibration curves were generated within the appropriate range for each of the incremental antibiotic influent concentration phases (i.e., 0.1-15 µg/L for 10 µg/L, 0.5-70 µg/L for 50 µg/L, and 1-300 µg/L for 250 µg/L). All calibration curve R^2^ values were > 99%. Both solvent-based and matrix-matched calibration curves were generated for all three compounds to ensure solvent based standards were representative of influent and effluent concentrations. Specifically, SMX, AMP, and ERY were dissolved in influent and effluent solutions at concentrations of 0.1, 1, 10, 100, and 1000 µg/L and processed using the same procedure employed to prepare reactor samples described above. Calibration curves of the results were plotted against standard solutions of the same concentrations dissolved in HPLC-grade water. For all three antibiotics, influent and effluent matrices showed equivalent concentrations and scaling at the ranges detected in the HPLC-grade water with R^2^ values of > 99.9. %. No isotope-labelled internal standards were used because the samples were collected and then analyzed by direct injection LC-MS on the same (no solid phase extraction required).

SMX, ERY and AMP were all targeted using positive ESI MS-Q-TOF mode. The LC gradient program for detection of all three compounds utilized 0.1% formic acid in water as mobile phase A and acetonitrile as mobile phase B as follows: t=0.0 min A=90% B=10%, t=3.0 min A=0% B=100%, t=5.0 min A=0% B=100%, t=5.10 min A=90% B=10%. LC conditions used included a flow rate of 0.4 mL/min, maximum pressure of 600 bar, column temperature of 40 ºC, and autosampler tray temperature of 8 ºC. A post-column switch was used to divert the first 0.5 min of column elution to waste to avoid sending hydrophilic compounds from the effluent matrix through the MS. Injection volumes ranged from 0.5-10 µL, depending on the target sample range for each operational phase, to ensure that no compound extracted ion chromatogram peaks exceeded saturation detection values. MS conditions used were as follows: sheath gas temp. of 400 ºC, sheath gas flow rate of 12 L/min, gas temperature of of 225 ºC, drying gas flow rate of 5 L/min, nebulizer pressure of 20 psi, capillary voltage of 3500V, nozzle voltage of 500V, acquisition rate of 1.5 spectra/s, and acquisition time of 667 ms/spectrum. Targeted compound acquisition parameters are provided in Table S2. All compound detection and quantification analyses were performed using the Agilent MassHunter Qualitative Analysis Navigator program.

**SI Table S3.** Targeted antibiotic properties and MS data acquisition parameters

| Compound | Molecular Weight (MW) | Retention Time (min) | MS Spectrum (m/z) | Fragmentor Voltage (V) |
| --- | --- | --- | --- | --- |
| Sulfamethoxazole | 253.052 | 2.17 | 254.059 | 400 |
| Erythromycin | 733.461 | 2.35 | 734.469 | 100 |
| Ampicillin | 349.110 | 1.44 | 350.117 | 400 |

### Quantification of ARGs using real time qPCR

All ARG standards used for qPCR quantification were constructed from PCR (Mastercycler nexus, Eppendorf, Hamburg, Germany) amplified products of DNA extracts from activated sludge (listed in Table S2). DNA samples from activated sludge. PCR products were analyzed on 2% agarose gel electrophoresis to verify the correct amplicon size. The correct DNA bands of interest were purified using Wizard^®^ SV Gel and PCR Clean-Up System (Promega, Madison, WI). Purified amplicons were cloned into linearized pMiniT 2.0 vector and transformed into NEB 10-beta Competent *E. coli* using the NEB PCR Cloning Kit (New England Biolabs, Ipswich, MA). Trasnformed E. *coli* were selected for on AMP selection plates. AMP resistant E. coli cells were harvested, and its plasmids was extracted using PureLink™ Quick Plasmid Miniprep kit (Invitrogen, Carlsbad, CA). Plasmid extracts were sent for Sanger sequencing (Laragen Sequencing & Genotyping, Culver City, CA) to confirm the presence of target ARG.. DNA concentration of the plasmid extracts were measured using the Quant-iT PicoGreen dsDNA Assay Kit (Invitrogen, Carlsbad, CA). Standard curves were established using serial dilutions of the purified plasmids (10^−1^ to 10^−8^). For all ARGs, the *intl*1 gene, and the *rpo*B gene, qPCR efficiencies ranged from 92% to 102%. Melting curve and gel electrophoresis were performed to assure the specificity of each qPCR reaction. SI Table 2 shows the forward and reverse primers, annealing temperatures, and thermal cycling conditions of all ARGs, *intl*1 and *rpo*B..

